# A single-cell genomic strategy for alternative transcript start sites identification

**DOI:** 10.1101/2021.12.09.472038

**Authors:** Yanling Peng, Qitong Huang, Rui Kamada, Keiko Ozato, Yubo Zhang, Jun Zhu

**Author notes:** **Corresponding authors**, Correspondence to Jun Zhu, or Yubo Zhang.

## Abstract

Alternative transcription start sites (TSSs) usage plays a critical role in gene transcription regulation in mammals. However, precisely identifying alternative TSSs remains challenging at the genome-wide level. Here, we report a single-cell genomic technology for alternative TSSs annotation and cell heterogeneity detection. In the method, we utilize Fluidigm C1 system to capture individual cells of interest, SMARTer cDNA synthesis kit to recover full-length cDNAs, then dual priming oligonucleotide system to specifically enrich TSSs for genomic analysis. We apply this method to a genome-wide study of alternative TSSs identification in two different IFN-β stimulated mouse embryonic fibroblasts (MEFs). We quantify the performance of our method and find it as accurate as other single cell methods for the detection of TSSs. Our data are also clearly discriminate two IFN-β stimulated MEFs. Moreover, our results indicate 82% expressed genes in these two cell types containing multiple TSSs, which is much higher than previous predictions based on CAGE (58%) or empirical determination (54%) in various cell types. This indicates that alternative TSSs are more pervasive than expected and implies our strategy could position them at an unprecedented sensitivity. It would be helpful for elucidating their biological insights in future.

## Main

Alternative transcription start sites (TSSs) usage is a common phenomenon and more than 50% genes contain multiple TSSs in human and mouse genomes ^1–4^. It plays an important role in gene transcription regulation in mammals ^5, 6^. For example, a novel transcript of the anaplastic lymphoma kinase (ALK) initiates from a *de novo* alternative transcription initiation site in ALK intron 19, which leads to the expression of a novel ALK isoform and results in melanomas ^5^. Similarly, alternative TSSs usage of tumor protein (TP) p53 family gene (p73) could result in full-length or shorter isoform. Elevated expression of the shorter isoform of p73 (deltaNp73), which inhibits apoptosis, has been associated with tumor progression and poor prognosis in several human cancers, including neuroblastoma, lung and ovarian carcinomas ^6^. Therefore, profiling of genome-wide alternative TSSs is expected to facilitate a better understanding of transcriptome complexity.

During the past decades, single-cell sequencing methods have made rapid advances on profiling genomics, transcriptomics, and epigenomics at unprecedented resolution ^7–9^. Traditionally, bulk cell sequencing methods can only reveal the average expression signal from an ensemble of cells assumed to be one homogeneous population/tissue status ^10^. Therefore, signals from small cell numbers or low densities could hardly be detected. Using single-cell technologies, numerous novel transcripts/cell types thereby have been identified ^11, 12^. For example, based on single-cell isoform RNA-Seq (ScISOr-Seq), 16,872 novel isoforms and 18,173 known isoforms in thousands of cerebellar cells have been detected ^12^. Based on single-cell universal poly(A)-independent RNA sequencing (SUPeR-seq), 913 novel linear transcripts and 2,891 circRNAs in mouse preimplantation embryos have been found ^11^. Furthermore, single-cell technologies are capable of dissecting heterogeneity in complex tissues ^13–15^. For example, single-cell methods enable classifying splenic cells into transcriptionally distinct groups ^13^ and identifying cell types of the central nervous system ^16^. Thus, single cell sequencing is able to detect cell heterogeneity and lowly expressed alternative TSSs which may be un-achievable by bulk cell technologies. To date, alternative TSSs annotation largely depends on computational predictions which have been available in Refseq ^1^, UCSC ^2^, and Genecode ^3^, as well as experimental identification by bulk cell technologies such as cap analysis of gene expression (CAGE) ^17^, oligo-capping ^18^, RNA annotation and mapping of promoters for the analysis of gene expression (RAMPAGE) ^19^. However, a paucity of single cell sequencing technologies are developed for alternative TSSs identification ^20, 21^.

Here, to enable genome-wide alternative TSSs annotation and cell heterogeneity detection, we have developed single cell TSSs sequencing (scTSS-seq) method (Figure 1). In this method, we first isolate single cells in Fluidigm C1 system, capture full-length cDNAs using SMARTer cDNA synthesis kit, and enrich 5’end cDNA tags specifically with customized dual priming oligonucleotides (DPO) ^22^ in Nextera XT DNA Library Prep Kit. DPO contains a longer 5’-segment, a shorter 3’segment, a poly(I) linker between these two segments ^22^. This enables enrichment of 5’end cDNA tags precisely even under sub-optimal PCR conditions ^22^. We then apply this method to naïve 1h and IFN-β pretreated 1h mouse embryonic fibroblasts (MEFs) (Materials and Methods) (Figure 2). Furthermore, in conjunction with RNA-seq, we examine whether pervasive alternative TSSs contribute to transcription memory following IFN-β restimulation. This study detects a larger proportion of genes containing alternative TSSs than empirical estimations, and finds that alternative TSSs are cell-type-specific and may be involved in transcriptional memory.

**Figure 1.**
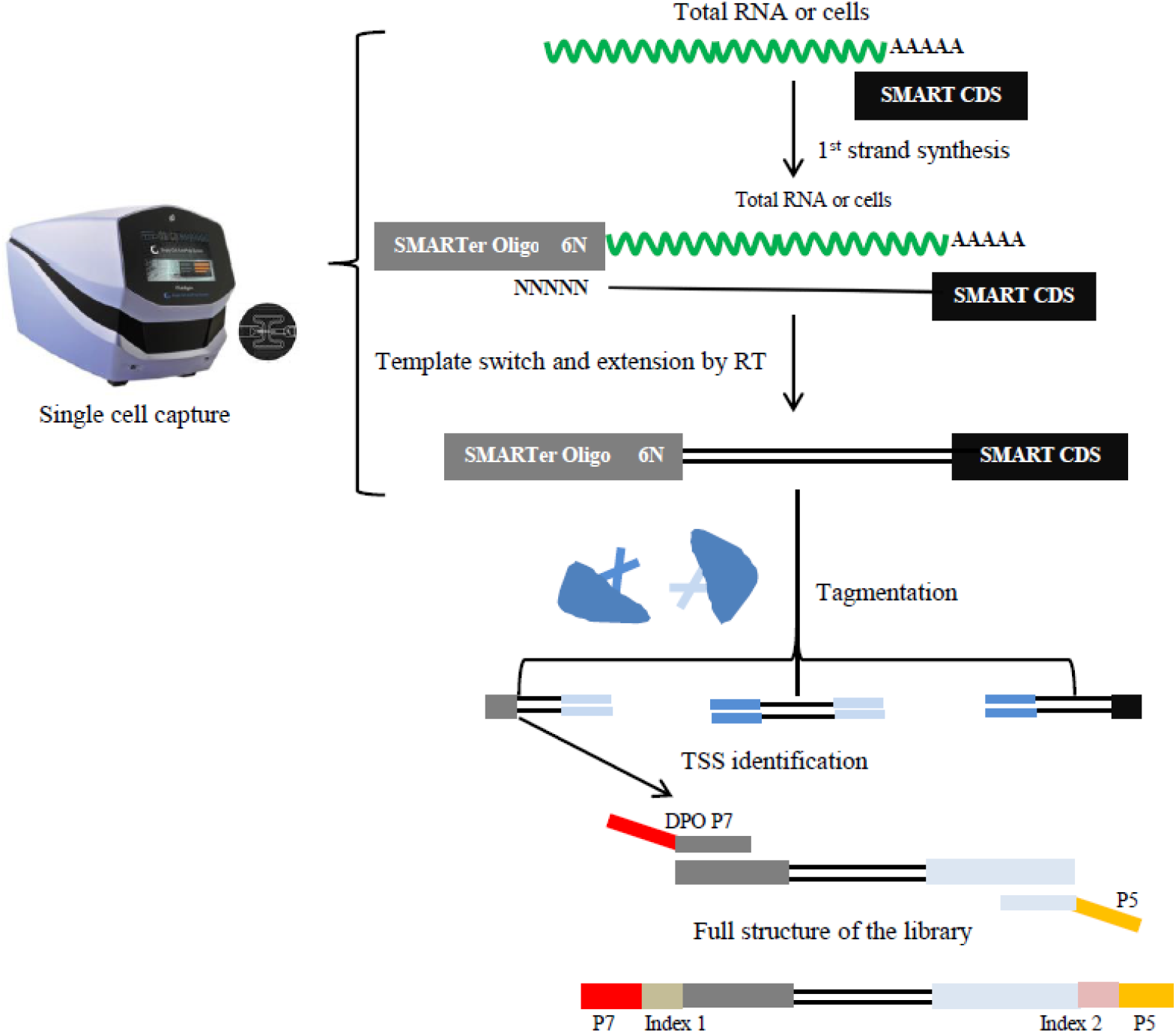
Schematic of the scTSS-seq method. P5, Nextera XT illumina P5 primers; DPO P7, custom-made dual priming oligonucleotides (DPO) P7 primers.

**Figure 2.**
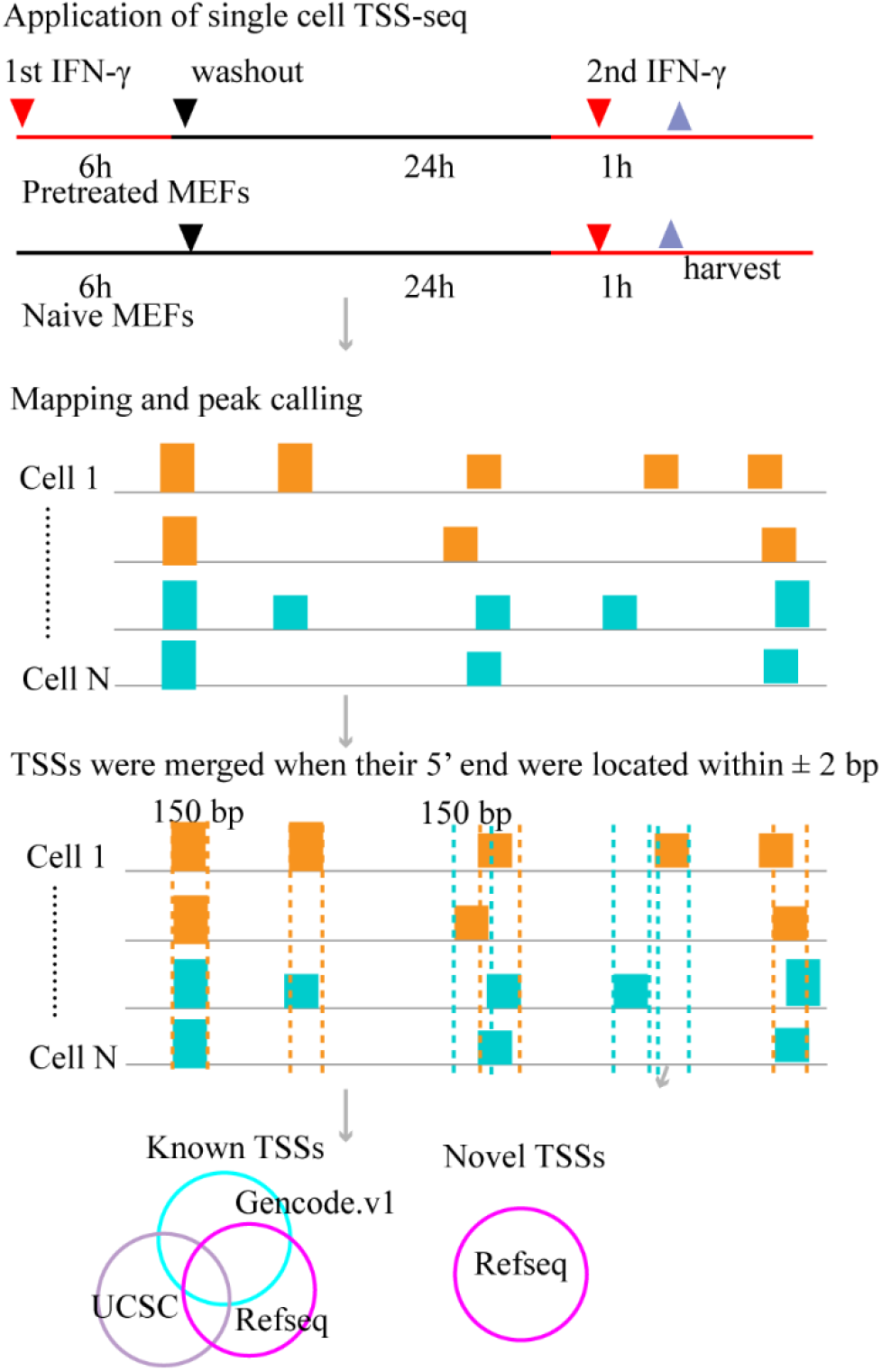
scTSS-seq applications and TSSs identification schematic in naïve 1h and pretreated 1h MEFs.

## Results

### The performance of scTSS-seq

Based on quality metrics, we removed six naïve and three pretreated cells (Materials and Methods). In the median, we detected 5,775 TSSs and 2,712 known TSSs (47% in total) per cell (Figure 3a). While, by applying C1 CAGE to 151 A514 cells following TGF-β stimulation (40 cells for 0h, 41 cells for 6h, and 70 cells for 24 h), Kouno et al. ^20^ detected a median of 2,788 CAGE clusters and 948 known TSSs (34% in total) per cell ^20^. When all single cells were pooled together, our study detected 518,097 TSSs in total and 143,923 known TSSs (28% in total). In comparison, Carninci et al. ^4^ detected 159,075 CAGE clusters in 145 different mouse libraries and 28,443 (18% in total) known TSSs using bulk-cell CAGE. Moreover, our results identified a median of 1,693 TSSs (30% in total) per cell and a total of 192,208 TSSs (37%) across all single cells overlapped with other TSSs in dbTSSs (Figure 3b), indicating a large part of TSSs that were not overlapped with annotated promoters were identified in other cell types. All the results suggested that our method was efficient at determining TSSs.

**Figure 3.**
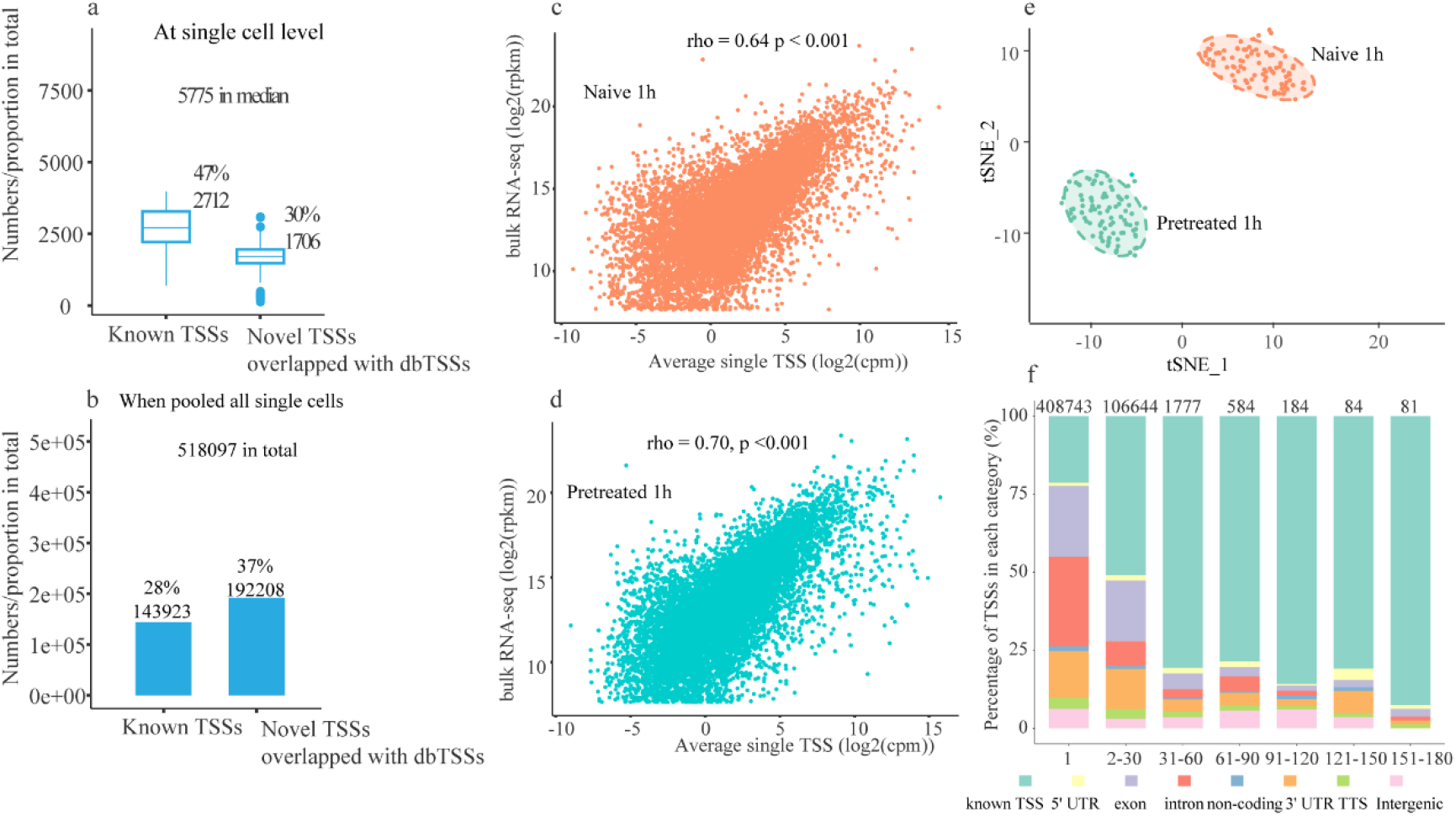
The performance of scTSS-seq. (a,b) TSSs identification at single cell level and when all single cells were pooled together. (c,d) The Spearman correlation of gene expression profile based on scTSS-seq data and bulk RNA-seq data for naïve 1h and pretreated 1h MEFs. (e) tSNE analysis based on TSSs expression levels. (f) Distribution of genomic annotation for TSSs. Note: 1, TSSs occurring in one cell; 2-30 TSSs occurring between 2 and 30 cells; 31-60, TSSs occurring between 31 and 60 cells; 61-90, TSSs occurring between 61 to 90 cells; 91-120, TSS occurring ranged from 91 to 120 cells; 121-150, TSSs occurring between 121 to 150 cells; 151-180, TSS occurring between 151 to 180 cells.

Later, we investigated the correlation of gene expression between scTSS-seq and bulk RNA-seq dataset for each cell type. For scTSS-seq data, gene expression levels were calculated as the sum of genic TSSs tags within the genes (cpm). The expression profiles of scTSS-seq data and bulk RNA-seq data were well correlated (rho = 0.64 for naïve 1h MEFs, rho=0.70 for pretreated 1h MEFs, *p* < 0.001, Spearman correlation, Figure 3c,d) compared to previous observations in other immune cells ^23 24^. This indicated that TSSs data were in good quality. To test whether scTSS-seq was able to differentiate cell types as well as other single-cell technologies, we employed the t-distributed Stochastic Neighbor Embedding (t-SNE) method for clustering. Two prominent clusters consistent with the two well-predefined cell types were classified (Figure 3e), suggesting that scTSS-seq could detect cell heterogeneity.

Furthermore, we characterized the distribution of TSSs across 183 high quality MEFs. About 408,743 TSSs were presented in one cell (cell-unique TSSs) while 109,354 occurred in more than one cell (cell-shared TSSs) (Figure 3f). About 25% of cell-unique TSSs were known TSSs, while more than 50% cell-shared TSSs were known TSSs (Figure 3f). These results further supported that single cell technologies were able to detect numerous novel TSSs or transcripts^11, 12^.

### Alternative TSSs detection at single cell level

To characterize the distribution of alternative TSSs and to compare with other bulk cell 5’end cDNA technologies, we first excluded the TSSs occurring in one cell and then counted alternative TSSs across naïve and pretreated MEFs. In this study, we found 109,354 TSSs occurring in more than two cells, of which 104,842 genic TSSs belonged to 11,761 Refseq genes. Among these genes, 9,712 (82% in total) had more than one TSS, and most genes contained 2~10 TSSs (Figure 4a). With bulk CAGE, 11,264 (58% in total) genes had alternative TSSs in mammals ^4^. High-density DNA microarrays had identified 1,609 (24% in total) of expressed genes having alternative TSSs in human fibroblasts cells ^25^. With empirical determination according to the UCSC database, 54% of these genes exhibited alternative TSSs ^26^. This suggested our method was capable of detecting alternative TSSs at genome-wide scale sensitively.

**Figure 4.**
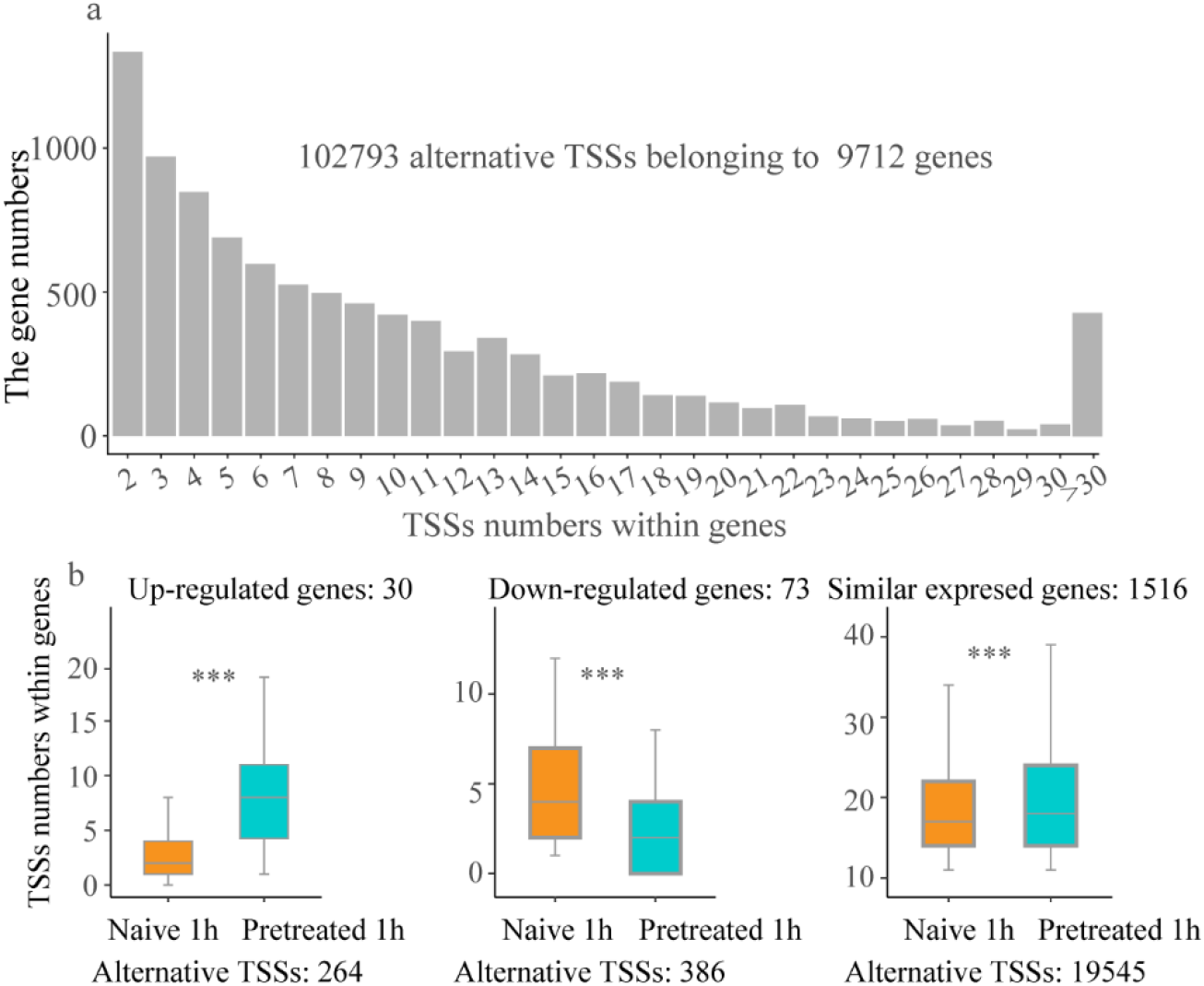
Alternative TSSs usage at single cell resolution. (a) The distribution of alternative TSSs. (b) The comparison of TSSs numbers within genes between naïve 1h and pretreated 1h MEFs for un-regulated, down-regulated and similar expressed genes. Note: ***, *p* < 0.001 the paired wilcoxon signed-rank test are performed.

To investigate whether alternative TSSs contributed to transcription regulation, we first assessed the alternative TSSs numbers for their significantly differentially expressed genes between two well-defined cell types. Based on single cell differential gene expression analysis, we detected 103 significant differentially expressed genes across two cell types, including 30 up-regulated and 73 down-regulated genes. The up-regulated genes included 264 alternative TSSs (Figure 4b). The TSSs numbers within these genes were significantly larger in pretreated 1h than in naïve 1h MEFs (*p* < 0.001, the paired wilcoxon signed-rank test with continuity correction). In contrast, down-regulated genes included 386 alternative TSSs. The TSSs numbers within these genes were significantly smaller in pretreated 1h than in naïve 1h MEFs (*p* < 0.001, the paired wilcoxon signed-rank test with continuity correction). Later, we investigated the changes of alternative TSSs numbers for similar expressed genes between two cell types. A total of 1,516 genes exhibited similar expression levels (fold change < 1.2, p value > 0.05), that included 19,545 TSSs. The numbers of alternative TSSs within these genes were significantly larger in pretreated 1h than naïve 1h MEFs (*p* < 0.001, paired wilcoxon signed-rank test with continuity correction). These results indicated that alternative TSSs may be involved in transcription regulation using varying mechanisms, such as dependent or independent of changes in gene expression.

### Alternative TSSs usage is correlated with transcriptional memory

With IFN-β re-stimulation in MEFs, it has been reported that the recruitment of RNA Pol II in memory IFN-stimulated genes (ISGs) (Materials and Methods) is faster and greater than that in non-memory ISGs ^27^. To test whether alternative TSSs usage affected transcription memory, we combined bulk RNA-seq data to analyze the changes of alternative TSSs numbers for ISGs across naïve and pretreated 1h MEFs. Taking similar strategies with Kamada et al. ^27^, we identified 112 up-regulated ISGs based on bulk RNA-seq data. These ISGs included 75 (67% in total) memory ISGs (*fold change* > 1.2 in pretreated 1h than naïve 1h MEFs) and 37 (33% in total) non-memory ISGs (*fold change* <= 1.2 and *fold change* > −1.2 in pretreated 1h than naïve 1h MEFs) (Figure 5a). GO analyses showed that memory and non-memory ISGs shared related categories, such as innate immune and defense responses. The GO results were similar in naïve and pretreated MEFs re-stimulated by IFN-β in previous study ^27^. Among these 75 memory ISGs, 910 alternative TSSs were detected (Figure 5b). The alternative TSSs numbers within memory ISGs were significantly larger in pretreated 1h than in naïve 1h MEFs (*p* < 0.001, the paired wilcoxon signed-rank test with continuity correction). For example, memory IGS *H2-K1* in pretreated 1h have more prevalent alternative TSSs usage than in naïve 1h MEFs (Figure 5c). A total of 226 alternative TSSs were identified belonging to 37 non-memory ISGs (Figure 5b). Alternative TSSs numbers within non-memory ISGs showed no significant differences between pretreated 1h and naïve 1h MEFs (*p* > 0.01, the paired wilcoxon test with continuity correction). These results supported that TSSs provided platform for RNA Pol II binding, while the rapid recruitments of RNA Pol II binding were necessary for transcription memory ^27, 28^.

**Figure 5.**
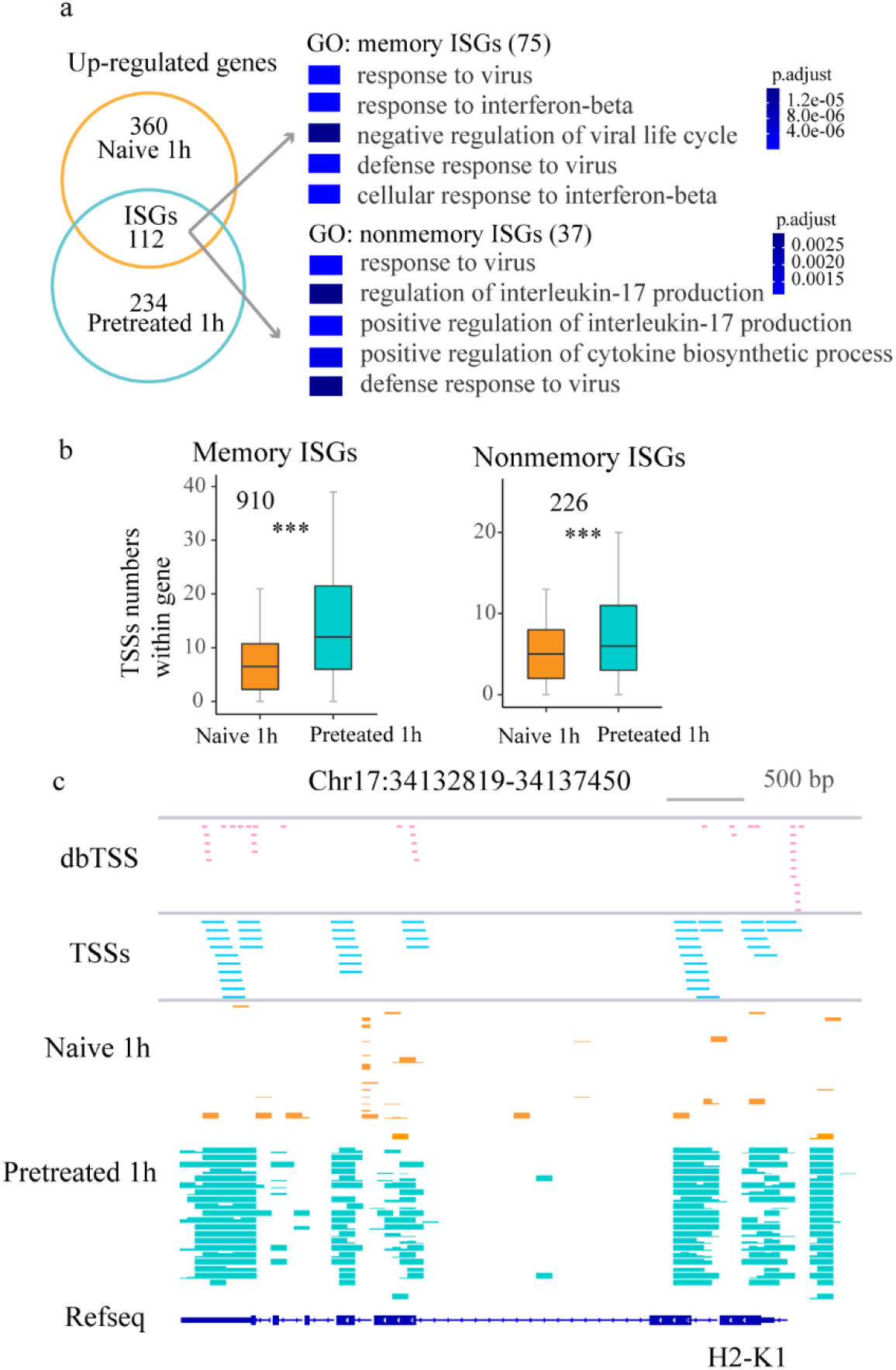
Memory ISGs related-TSSs biological process. (a) The identification of memory and non-memory ISGs and GO analysis for the identified memory and nonmemory ISGs. (b) Comparison of alternative TSSs numbers per gene between naïve 1h and pretreated 1h MEFs, the paired wilcoxon signed-rank test are performed, (c) Example of TSSs distribution for ISGs, *H2-K1*.

## Discussion

In this study, we have developed scTSS-seq for genome-wide alternative TSSs detection at single cell level. This method utilizes Fluidigm C1 system to increase detection sensitivity of transcripts and genes ^29^, SMART technology to increase full-length rate of cDNAs ^30^ and DPO system to increase enrichment specificity ^22^. Meanwhile, the results show that the scTSS-seq method is efficient at determining TSSs compared to C1 CAGE ^20^. In addition, this method is in good performance by estimating the correlation between single cell TSSs and bulk RNA-seq expression profile. Furthermore, higher rates (82%) of alternative TSSs across naïve 1h and pretreated 1h MEFs are detected in comparison with previous predictions based on CAGE (58%) or empirical determination (54%) in various cells. Thus, scTSS-seq is an efficient method for positioning alternative TSSs at genome-wide scale.

At the same time, we have detected that memory ISGs have larger numbers of alternative TSSs in pretreated 1h than in naïve 1h MEFs, implying that alternative TSSs usage are cell-type-specific and may be contributed to transcriptional memory ^27^. However, it still remains challenging to validate the role of alternative TSSs on memory ISGs in single cells so far. Firstly, cell viability is required when isolating single-cells for the purpose of production of monoclonal cell culture ^31, 32,^ which would be affected during single cell separation ^32, 33^ Secondly, the amount of protein and RNA of single cells may be not enough for histochemical experiments ^34, 35^. Thirdly, the quality and amount of antibodies would be limited for validation ^36, 37^. Despite these factors, we develop a reliable method for genome-wide alternative TSSs detection and also dissect a more pervasive alternative TSSs phenomenon in cells. Meanwhile, we provide useful alternative TSSs resources for future biological functional research.

## MATERIALS AND METHODS

### Cell preparation

The preparation of MEFs was similar to Kamada et al ^27^. In brief, MEFs were separated from day 13.5 mouse embryos, and then cultured in Dulbecco’s minimal essential medium (DMEM) with 10% fetal bovine serum (FBS) at 37 °C with 5% CO_2_. Later, they were treated with 100 units/mL of murine recombinant IFN-β (PBL Interferon Source) with two different ways for indicated periods (Figure 2). On the one hand, MEFs were treated without IFN-β for 6h, washed, and left without IFN-β for 24h (naïve 0h MEFs). On the other hand, MEFs were treated with IFN-β for 6h, washed, and then left without IFN-β for 24h (pretreated 0h MEFs). In this study, both naïve and pretreated MEFs were re-stimulated with IFN-β for 1 hour (naïve 1h and pretreated 1h MEFs) and harvested for scTSS-seq (Figure 2). For bulk cell RNA-seq, besides naïve 1h and pretreated 1h MEFs, naïve 0h and pretreated 0h MEFs were harvested.

### scTSS-seq experiment procedure

96 naïve 1h and 96 pretreated 1h MEFs were captured using the C1 Fluidigm system according to the manufacturer’s instructions. Upon capture, reverse transcription and cDNA preamplification were performed in the C1 Fluidigm system using the SMARTer PCR cDNA Synthesis Kit (Clontech 634926) according to manufacturer’s recommendations. Full-length cDNAs were harvested and diluted to a range of 0.2 ng/ul, and Nextera libraries were prepared using the Nextera DNA Sample Preparation Kit (Catalog No. FC-131-1096). A Nextera P5 (Illumina) and a custom-made P7 index primer (Supplementary Table S1) were used to amplify the tagmented fragments. The P7 index primers were designed as DPO systems. The details of the scTSS-seq library construction procedure were given in supplementary file 1. Libraries were pooled and single-end sequencing was performed on HiSeq 2000 (Illumina).

### scTSS data processing

Reads were subjected to adapter and quality trimming with Trim Galore (https://www.bioinformatics.babraham.ac.uk/projects/trim_galore/; v0.4.4; options: default parameters). The trimmed reads were then aligned to the mouse genome (mm9) using bowtie2 ^38^ (v2.29, options: default parameters). HOMER ^39^ tag directory was created with ‘makeTagDirectory’ function (options: default parameters) for each cell. For each tag directory, peaks were called with HOMER ‘findpeaks’ function ^39^ (options: -style tss -ntagThreshold 200). For each cell, peaks were then annotated with HOMER ‘annotatePeaks.pl’ function ^39^, three different mm9 transcript models were used for annotations, including Refseq (HOMER default), UCSC (https://genome.ucsc.edu/cgi-bin/hgTables), and Genecode V1 (https://www.gencodegenes.org/mouse/release_M1.html). In these models, annotated promoters were defined as regions between downstream 1000 bp and upstream 100 bp of TSSs. For each cell, peaks overlapped with annotated promoters in any of three models were termed as known TSSs, while the remaining were novel TSSs.

Low quality cells would have lost transcripts prior to cell lysis and their exon numbers would be larger in broken cells ^40^. Thus, we used the ratios between known TSSs numbers and identified TSSs numbers in total for each cell (known TSS ratios) for filtering low quality cells. In this study, we used threshold as z-score of known TSSs ratios <= −1 for removing cells. The remaining cells were considered as high quality cells and used for downstream analysis.

To produce the matrix of TSSs tags across cells, we merged TSSs when their 5’ end were located around ±2bp across all high quality cells. For further analysis, the expression levels of TSSs were normalized to counts per million, with calculation by counts per mapped reads per 10 million. The expression levels of genes were calculated as the expression levels of TSSs tags within Refseq genes on average. TSSs clustering and differential analysis were performed by using R packages Seurat ^41^. To know how many novel TSSs were overlapped in other TSSs datasets across difference cell types, we downloaded TSSs coordinates (bed format) from different cell types or tissues such as 10Thalf, 3T3, ATDC5 and embryos (ftp://ftp.hgc.jp/pub/hgc/db/dbtss/dbtss_ver8/mm9/TSSseq/bed/).

### RNA-seq procedure and analysis

The library construction of RNA-seq was similar to Kamada et al. ^27^. In contrast, single-end sequencing on a HiSeq2000 (Illumina) was used in our study. The obtained sequences were aligned to the mouse genome mm9 using Tophat ^42^. Transcript abundance was quantified using Cufflinks 1.2.1 ^43^. Genes showing significantly (*p* value < 0.01) higher transcript expression (RPKM) in treated cells following IFN treatment (1h) over untreated cells (0h) were considered up-regulated genes, termed as IFN stimulated genes (ISGs) and were analyzed further. GO analysis of different categories of ISGs were performed by using R packages ClusterProfiler ^44^, respectively. Terms with *p* < 0.05 were regarded as significant enrichment.

## Supporting information

Information of custom-made DPO P7 primers

The details of scTSS-seq library construction procedure

## Data availability

scTSS-seq and RNA-seq raw data has been submitted to the Gene Expression Omnibus (GEO) under accession GSE174480.

## Supplementary

Table S1 Information of custom-made DPO P7 primers

File 1 The details of scTSS-seq library construction procedure

## Acknowledgements

This work has been supported by National Key Research and Development Program of China [2018YFA0903201], National Natural Science Foundation of China (grant no. 2017M620977), the Science, Technology and Innovation Commission of Shenzhen Municipality (grant no. JCYJ20180306173714935).

## Contributions

Y.Z. and J.Z.conceived the project, supervised, and designed the study. Y.Z., and R.K. performed experiments. K.O. supervised the culture study. Y.P. and Q.H. carried out computational analyses. Y.P., Y.Z. and J.Z. wrote the manuscript.

