## Supplementary material for "A single-cell genomic strategy for alternative transcript start sites identification": The details of scTSS-seq library construction procedure

Single cell TSS experiment (96 wells, combined it with Nextera XT library construction)

1. Measure the concentration of single cell samples (with PicoGreen or GX), normalize the sample to ~0.1-1 ng for Nextera XT
2. Take 48 ~0.5 ng cDNA from each C1 pretreated_1h and Naïve_1h plates and put them into a new 96 well plate, add water into each well to make the volume to 2.5 µl each (0.2 ng/µl).
3. Add 5 µl of TD Buffer (Nextera XT kit, ensure the buffer is fully dissolved) to each well and change tips between samples.
4. Add 2.55 µl of ATM to the wells containing input DNA and TD Buffer. Gently pipette up and down 5 times to mix.
5. Centrifuge at 280xg at 20°C for 1 minute.
6. Place the NTA plate in a thermocycler and run the following program:
   • 58°C for 10 minutes
   • Hold at 10°C
7. Once the sample reaches 10°C proceed immediately to Neutralize NTA. Carefully remove the Microseal "B" seal and add 2.5 µl of NT Buffer to each well of the NTA plate (total volume should be 12.5 µl now).
8. Split the sample into two equal parts, one for single cell TSS experiment (following part) and one for storage

Single cell TSS experiment:

1). Add 7.5 µl Ampure beads (1.2:1) to each well. Gently pipette up and down 5~10 times to mix.

2). Incubate at room temperature for 5 mins. Place beads on magnetic stand for 5min or the solution is cleared;

3). Remove supernatant and wash the beads twice with 100 µl **FRESHLY** made 80% EtOH, which is added while bead is on magnetic stand and incubate at RT for 30'' before removing the liquid.

4). After the 2nd wash, briefly spin the tube, put back to magnetic stand, remove the residue EtOH with 10 µl pipette, and dry @ RT for 5mins;

5). Add 16 µl water and mix well by pipetting or brief vortex. Put the tube back to magnetic stand. Wait the solution becomes clear or 5mins.

6). Add primers to a fresh plate with the following barcode (96 samples):

| | |
|---|---|
| P7_SMDP (5 uM, N701-N712) | 1.5 ul |
| P5 (5 uM, XT kit, N501-N508) | 1.5 ul |

7). Transfer the 15 µl supernatant to the primer plate.

8). Prepare 100× PCR reaction mix.

| | |
|---|---|
| dNTP (100mM) | 0.6 ul |
| Mg2+ (25mM) | 1.5 ul |
| Buffer 10× | 3 ul |
| pfu CX | 0.6 ul |
| H2O | 6.3 ul |

9). Distribute the 12 µl PCR reaction mix to each well of tagmentated cDNA plate. Mix well and briefly spin.

10). Run following PCR program for PA site enrichment.

| STEP | Tag Pol. |
| --- | --- |
| Initial Denaturation | 95°C (3mins) |
| 20 Cycles | 95°C (15s) |
| | 54°C (15s) |
| | 72°C (30s) |
| Final Extension | 72°C (5mins) |
| Hold | 10°C |

11). Take out 5 µl PCR product to run gel for QC (**select 5 highest and 5 lowest concentration samples according to the original C1 plates for QC**).

12). Take 2 µl each sample of PCR products and pool them together.

13). Take 20 µl pooled sample to run 2% agarose gel to do size selection (**200bp–300bp**). Use the gel purification kit to pure the gel PCR product.

**a. Excise the DNA fragment from the agarose gel using a razor blade, transfer it into a 1.5 ml microcentrifuge tube.**

**b. Add 3 volumes of ADB (~600 µl) to each volume of agarose excised gel.**

**c. Incubate at 55 °C for 10 minutes until the gel slice is completely dissolved.**

**d. Transfer the melted agarose solution to a Zymo-Spin™ Column in a Collection Tube. Centrifuge for 30s with max speed. Discard the flow-through.**

**e. Add 200 µl of DNA Wash Buffer to the column and centrifuge for 30s with max speed. Discard the flow-through. Repeat the wash step.**

**f. Add 20 µl DNA water directly to the column matrix. Incubate 1 min. Centrifuge for 60s with max speed to elute DNA.**

14). Quality the DNA with Quibit. And the library would be ready for sequencing.
